# PEG400 and Tween 80–based simple vehicle for oral delivery of peptide compounds in fish

**DOI:** 10.1101/2025.09.03.673866

**Authors:** Sayaka Matsuo, Takehiko Yokoyama, Shun Satoh, Yuki Yoshio, Kazuya Fukuda

## Abstract

Pharmacological approaches are increasingly used in diverse fields of fish biology. However, injection-based administration can induce stress or injury, potentially interfering with the natural physiological and behavioral states of the fish. Here, we present a non-invasive vehicle for oral administration of peptide-based compounds. To demonstrate its utility, the oxytocin receptor antagonist atosiban was administered orally using this vehicle. Atosiban was then qualitatively detected in serum by high-performance liquid chromatography. This study provides a proof of concept for an easy-to-use and minimally disruptive tool for various fish experiments.

## Introduction

In recent years, studies of fish behavior and other areas of fish biology have increasingly adopted integrative approaches that consider not only ultimate factors but also proximate mechanisms (Davies et al. 2012; Rubenstein and Alcock 2019). This broader perspective has enabled researchers to gain a more comprehensive understanding of the evolution of behavior and to identify conserved patterns across taxa. It has also led to novel insights that are difficult to obtain through observational studies alone. For example, Matsumoto et al. (2018) investigated filial cannibalism in a blenniid fish, *Rhabdoblennius nitidus*, and resolved a theoretical paradox in behavioral ecology by incorporating an endocrinological approach. Similarly, Nowicki et al. (2020) compared expression patterns of nonapeptides, dopamine, and opioid receptor genes in the forebrains of pair-bonded and solitary butterflyfish species and found that these systems modulate sociality, as also observed in other vertebrates. These studies illustrate that approaches bridging proximate and ultimate factors can open new directions for research in fish biology.

An accessible approach to bridge these levels is pharmacological manipulation, in which candidate regulatory systems (e.g., neural, endocrine, or physiological bases) are enhanced or inhibited to assess their functional consequences. This approach has been widely used, for example, in studies of territorial behavior (Santangelo and Bass 2006), paternal care (O’Connell et al. 2012), shoaling (Ataei Mehr et al. 2020), and reproductive behavior (Almeida et al. 2023). While genetic tools, such as genome editing, are increasingly available, they remain limited in many non-model fish species. Given the great diversity of fishes, pharmacological manipulation will continue to be an important and broadly applicable approach. In many cases, pharmacological manipulations involve administering compounds via intraperitoneal, intracerebroventricular, or intramuscular routes. However, because such injection-based methods are inherently invasive (e.g., Håstein et al. 2005), they can be unsuitable depending on the experimental design. For example, when experiments must be conducted over extended periods, treatments are required to be administered repeatedly, which may induce stress responses and/or physical injury, thereby altering physiological state and potentially confounding behavior. To avoid these problems, non-invasive compound delivery methods are needed.

One non-invasive method is oral administration, which has been used in long-term treatments (e.g., Madrid et al. 2021; Lu and Patton 2022; Ochocki and Kenney 2023). However, oral administration of many compounds remains challenging compared with injection-based methods because of physiological and biochemical barriers, including low permeability and degradation in gastrointestinal fluids (Song et al. 2004; Lou et al. 2023). In fish, oral administration faces similar limitations, and efforts have been made to improve its efficacy (e.g., Amezawa et al. 2018). A practical and accessible strategy is to optimize the composition of the vehicle mixed with compounds to enhance delivery efficiency, as vehicle composition is known to influence oral delivery outcomes (e.g., Tønsberg et al. 2010; Serdoz et al. 2010). Establishing an effective and easy-to-use vehicle system could provide a valuable tool for experimental studies across diverse fields of fish biology.

In this study, we examined whether an oligopeptide compound could enter systemic circulation following oral administration in combination with a vehicle modified with reference to pharmaceutical research. As a model fish, we used the convict cichlid (*Amatitlania nigrofasciata*), a species employed in various behavioral studies, including those on reproductive behavior (e.g., Santangelo 2005), proactive prosociality (Satoh et al. 2021), and affective state (Laubu et al. 2019). As a test case, we selected atosiban, an oxytocin receptor antagonist that has been widely studied in the context of social behavior. In some fish, atosiban introduced into systemic circulation via intramuscular or intraperitoneal injection affects behavior, with its effects expected to result from actions within the central nervous system (e.g., Cardoso et al. 2015; Ataei Mehr et al. 2020). However, to our knowledge, it has not yet been used via oral administration in teleost fish. Therefore, we used high-performance liquid chromatography (HPLC) to determine whether orally administered atosiban, given with the modified vehicle, can be detected in serum. Based on these results, we propose a vehicle-based oral administration approach as a minimally disruptive option for pharmacological studies in fish.

## Materials and methods

Six male convict cichlids (58–99 mm SL) were collected from Okinawa-jima Island, Japan. The fish were housed in a 50 L tank at Kitasato University, where the water temperature was maintained at 26°C under a 14 h light/10 h dark photoperiod, and were fed a commercial fish food (Hikari Carnivore, Kyorin Co., Ltd.; hereafter, “food”) twice daily.

To prepare the medicated food for oral administration, atosiban (T5148, TargetMol) was first dissolved in dimethyl sulfoxide (DMSO; 045-24511, Fujifilm Wako Pure Chemical Co.) and then mixed to prepare either a modified formulation or a control solution. The modified formulation was prepared following Kim et al. (2017) and consisted of atosiban (50 µg/µl), DMSO (5% [v/v]), polyethylene glycol 400 (PEG400; N0443, Tokyo Chemical Industry Co., Ltd.) (40% [v/v]), Tween 80 (35703-62, Nacalai Tesque, Inc.) (5% [v/v]), and reverse osmosis water. The control solution contained atosiban (50 µg/µl) and DMSO (5% [v/v]) in reverse osmosis water to control for the absence of the other excipients. Food pellets were cut into pieces approximately 3 mm in diameter and 4 mm in length, and each piece was infused with 10 µl of either the modified formulation or the control solution, then air-dried at room temperature for 1 hour. All medicated food was prepared immediately before oral administration.

The procedures for oral administration and sample collection were as follows. After a 3-day fasting period, the fish were divided into two groups. Prior to the experiment, all fish underwent a 1-to 2-week training period. During this period, the fish were fed twice daily at fixed times to ensure that they would eat the food immediately after it was introduced into the tank. In the experiment, three individuals were each fed three pieces of food treated with the modified formulation, and the remaining three were each fed three pieces of food treated with the control solution. Consistent with the training period, all individuals consumed the treated food immediately (within 30 sec of introduction). After 55 min, each fish was deeply anesthetized with benzocaine, and blood was collected from the caudal vessel. The fish were euthanized as a terminal procedure following blood collection. Blood samples were immediately centrifuged at 10,000 × g for 15 min at 4°C to obtain serum. As preliminary trials indicated that the serum volume obtainable from a single individual was insufficient for HPLC analysis, the sera from the three fish that had been fed the same treated food were pooled. After the addition of 50 µl of 1 M perchloric acid (162-00695, Fujifilm Wako Pure Chemical Co.) to the pooled sample, the sample was centrifuged again at 15,000 × g for 20 min at 4°C, and the supernatant was collected. The supernatant was then passed through a 0.22 µm hydrophilic nylon filter (SLNY1322N, Hawach Scientific) and concentrated to approximately 50 µl using a centrifugal concentrator. The concentrated samples were then injected into the HPLC system.

Chromatographic analysis was performed with reference to Raju et al. (2014) using a JASCO (Tokyo, Japan) HPLC system consisting of a PU-4180 quaternary gradient pump with a degasser, a CO-4060 column oven, an AS-4050 autosampler with a cooling system, a UV-4075 UV/VIS detector, and a ChromNAV data processor. The analytical column was a reversed-phase ODS-120H (4.6 × 250 mm; Tosoh, 0023415). Elution was performed with a mixture of solvent A (30 mM potassium phosphate buffer at pH 3.2) and solvent B (acetonitrile:Milli-Q water = 80:20) at 40°C; the flow rate was set at 1.0 ml·min^-1^ and the detection wavelength was set at 220 nm. The elution gradient was set as follows: 0–5 min, 20–20% solvent B in solvent A; 5–12 min, 20–70% solvent B in solvent A. Because the primary objective of this study was to qualitatively examine whether atosiban could be detected in systemic circulation as a proof of concept, rather than to determine precise pharmacokinetics, a calibration curve for quantification was not generated, as the method was not validated for quantitative recovery or matrix effects in serum. Standard solutions of atosiban were prepared at 0.5 and 5 µg/mL in ultrapure water and analyzed under the same chromatographic conditions. Both standards were used to confirm retention time, and the chromatogram shown for each group uses the standard concentration that provided the clearest visualization relative to the background signal in the corresponding serum chromatogram.

## Results

HPLC analysis detected a peak corresponding to atosiban, exhibiting the same retention time as the standard, only in the pooled serum sample from the group that ingested food treated with the modified formulation (Fig. 1). No such peak was observed in the group fed the control solution. These results are consistent with the presence of atosiban in serum after oral administration using the modified formulation.

**Fig 1.**
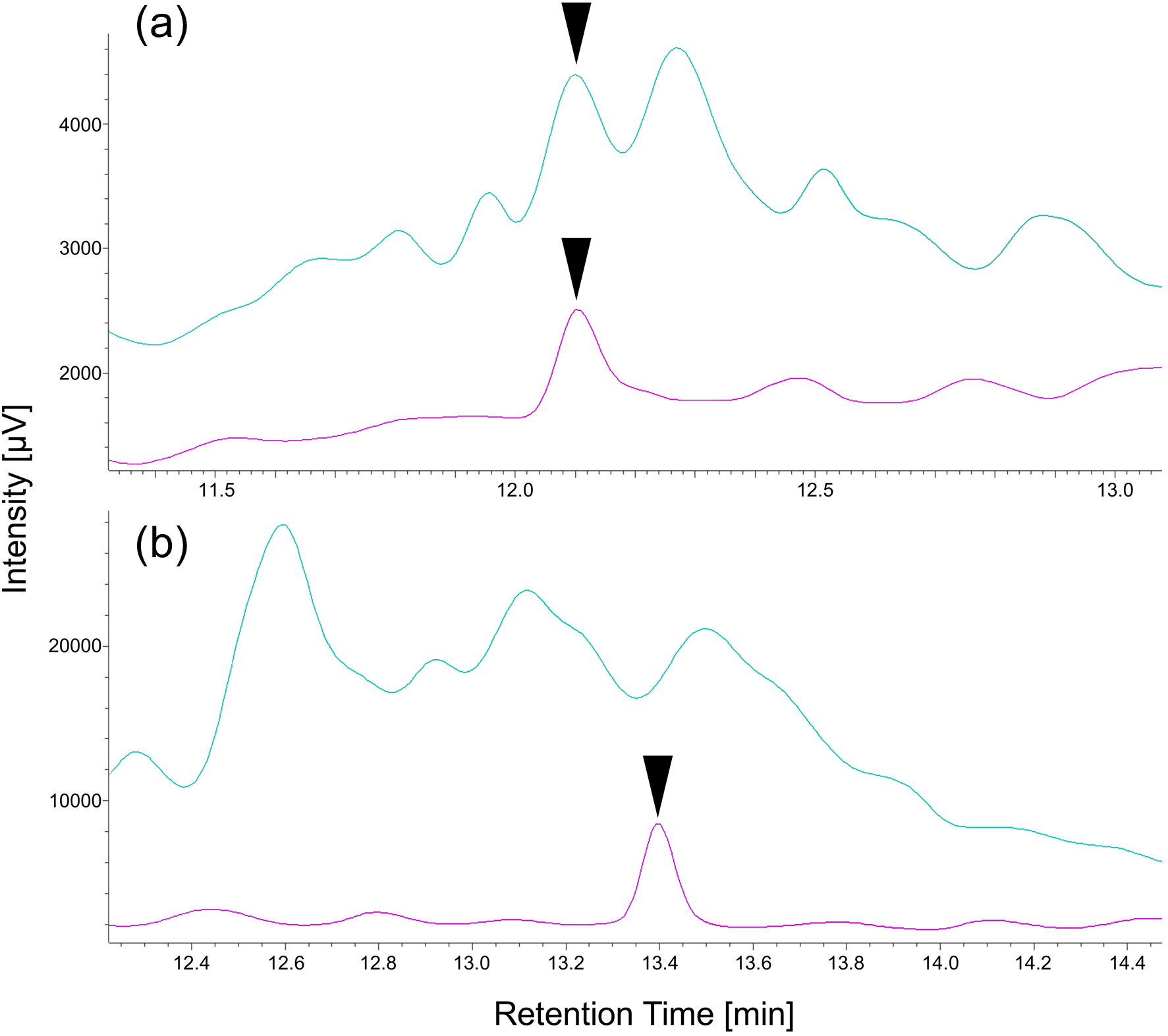
Chromatograms of (**a**) a blood extract from fish that ingested atosiban with the modified vehicle solution, and of the 0.5 µg/ml atosiban standard, and (**b**) a blood extract from fish that ingested atosiban with the control solution, and of the 5 µg/ml atosiban standard. Black arrowheads indicate the peak of atosiban. Cyan and magenta lines represent the chromatogram of the blood extract and standard, respectively.

## Discussion

Ethological studies based on adaptationist thinking have long contributed to understanding the evolutionary background (ultimate factors) of social behaviors across vertebrates, and some theoretical and empirical frameworks of ethological studies are broadly applicable to adjacent research fields (Davies et al. 2012; Rubenstein and Alcock 2019). One of the key emerging questions is whether behaviors that evolved under similar ultimate factors are underpinned by entirely different phylogenetic origins and mechanisms, or whether they are, at least in part, supported by evolutionarily conserved mechanisms. Indeed, commonalities across vertebrates have been identified not only at the phenotypic level but also in regulatory mechanisms underlying some social behaviors, such as territoriality (Oldfield et al. 2015), parental behaviors (Dulac et al. 2014), and monogamous mating systems (Young et al. 2019). These examples highlight how integrative approaches can reveal conserved proximate mechanisms in fish and other vertebrates. In this context, the vehicle proposed here provides a practical option for incorporating minimally invasive pharmacological manipulation into fish experiments without complex procedures or costly equipment.

PEG400 and Tween 80 are excipients used for oral administration and are generally regarded as nontoxic and nonirritant in mammals (e.g., Rowe et al. 2009; Gullapalli and Mazzitelli 2015; Ma et al. 2017). In rats, PEG400/polysorbate 80 formulations improved the oral bioavailability of the poorly water-soluble drug halofantrine, likely through formulation-dependent effects on solubility and precipitation upon dilution (Tønsberg et al. 2010). Although the mechanism in the present study remains unresolved, the detection of atosiban in serum after oral administration may likewise reflect formulation-dependent effects within the gastrointestinal tract. Related formulations have also been applied in fish (e.g., Kim et al. 2017). We expect that the vehicle proposed here will be particularly useful for experiments requiring minimally disruptive conditions, including long-term studies or repeated administrations.

On the other hand, several considerations should be noted in applying our method. First, although we used atosiban as a model compound, the extent to which this vehicle improves oral delivery efficiency for other substances should be evaluated individually. For example, while atosiban is an oligopeptide and hydrophilic, improved oral delivery with PEG400/polysorbate 80 formulations has been reported for a poorly water-soluble hydrophobic drug (e.g., Tønsberg et al. 2010). Second, if greater transfer into systemic circulation via the oral route is required, alternative strategies may be necessary. In pharmacology, several sophisticated approaches for improving oral bioavailability have been developed (Brayden et al. 2020; Cui et al. 2022; Lou et al. 2023). As our primary aim was to minimize procedural complexity, such methods would be worth considering if improving bioavailability is prioritized over simplicity. Therefore, the present findings should be regarded as a preliminary proof of concept. Elucidating the underlying mechanisms was beyond the scope of this study and will require future research. Nevertheless, we hope that our results will encourage the future development of non-invasive pharmacological methods for application in experimental studies of fish biology.

## Acknowledgments

We are grateful to C. Ogawa for their support during our study. This work was financially supported by JSPS KAKENHI Grant Numbers 24932659, 20346357, 23721203, and the Hakubi Project of Kyoto University.

## Author contributions

Conceptualization: Kazuya Fukuda; Methodology: Sayaka Matsuo, Takehiko Yokoyama, Shun Satoh, Yuki Yoshio, Kazuya Fukuda; Formal analysis and Investigation: Sayaka Matsuo, Takehiko Yokoyama, Kazuya Fukuda; Writing - original draft preparation: Sayaka Matsuo, Kazuya Fukuda; Writing - review and editing: Takehiko Yokoyama, Shun Satoh, Kazuya Fukuda; Funding acquisition: Shun Satoh, Kazuya Fukuda; Resources: Sayaka Matsuo, Shun Satoh, Kazuya Fukuda; Supervision: Takehiko Yokoyama, Shun Satoh, Kazuya Fukuda

## Declarations

### Ethical approval

At Kitasato University, there are currently no institutional ethical regulations specifically for research on fish. Therefore, all procedures were conducted in accordance with the Guidelines for the ethical treatment of nonhuman animals in behavioral research and teaching of the Association for the Study of Animal Behaviour and Animal Behavior Society (ASAB Ethical Committee/ABS Animal Care Committee, 2023), as well as the Guidelines for the use of fishes in research published by the Ichthyological Society of Japan in 2003 (https://www.fish-isj.jp/message/guideline/2003_1011/).

The convict cichlid was introduced into Japan as an ornamental fish, and individuals that subsequently entered natural waters are thought to have established a feral population on Okinawa-jima Island sometime between approximately 1970 (Tachihara *et al*., 2002) and 1989 (Ishikawa *et al*., 2013). All specimens used in this study were collected from this feral Okinawa-jima population. This study complied with Japanese law and the guidelines of the Ichthyological Society of Japan.

